# Conserved water molecules as structural ligands modulating pathogenic variation in human protein binding sites

**DOI:** 10.64898/2026.04.01.715522

**Authors:** Janez Konc, Karmen Recer, Tanja Kunej, Dušanka Janežič

## Abstract

Conserved water molecules (CWMs) are tightly bound solvent molecules that occupy well-defined and recurrent positions in protein structures. Although they are known to influence protein stability, function, and ligand binding, their contribution to human genetic disease has remained largely unexplored. Here, we demonstrate that CWMs substantially contribute to the pathogenicity of single nucleotide polymorphisms (SNPs). By systematically mapping SNPs onto ligand-binding and conserved water sites across human protein structures in the Protein Data Bank, we find that pathogenic variants are strongly enriched at CWM positions. Enrichment is particularly pronounced at CWM sites within ligand-binding regions, exceeding that observed for ligand-binding sites as a whole. To establish a mechanistic link, we performed molecular dynamics simulations on human lysosomal acid glucosylceramidase (GCase), encoded by GBA1 and associated with Gaucher disease and Parkinson’s disease risk. Removal of a single conserved water molecule in the wild-type protein recapitulates key structural features of the pathogenic L444P variant, whereas stabilization of this water in the mutant restores native-like behavior. These findings demonstrate that disruption of a conserved water molecule can induce long-range structural changes consistent with disease-associated mutations. Together, our results identify conserved water molecules as functional structural elements whose disruption represents a recurrent mechanism of protein dysfunction and provide direct mechanistic evidence for their pathogenic role in Gaucher disease.

**Significance Statement:** Are conserved water molecules hidden determinants of human genetic disease? By systematically mapping single nucleotide polymorphisms (SNPs) onto protein structures, we show that pathogenic variants are strongly enriched at conserved water positions, particularly within ligand-binding sites. This enrichment exceeds that observed for most previously studied ligand types, including small molecules, nucleic acids, and proteins. Molecular dynamics simulations further reveal that disruption of a single conserved water molecule can induce long-range structural effects consistent with disease-associated variants in lysosomal acid glucosylceramidase, linking these findings to Gaucher disease and Parkinson’s disease risk. Together, our results establish conserved water molecules as previously underappreciated structural determinants of human disease and highlight their relevance for understanding genetic variation and guiding drug discovery.

## Introduction

Proteins interact through binding sites with diverse ligands, including other proteins, nucleic acids, metal ions, cofactors, small molecules and conserved water molecules (CWMs). Among these, CWMs remain underexplored, despite mounting evidence of their essential roles in protein fold stability, ligand recognition, and fine-tuning of protein dynamics (1–6).

Single nucleotide polymorphisms (SNPs) that perturb binding sites can disrupt protein function by altering ligand recognition, or by causing protein destabilization or aggregation, often driving disease (7–10). Structural context offers a powerful framework for predicting such effects (11, 12), aided by the growing coverage of human structures in the Protein Data Bank (PDB) (13) and high-accuracy models (14, 15). Proteome-scale resources now integrate these structures for variant pathogenicity predictions (16), while genomic repositories (17, 18) currently catalog hundreds of millions of variants from large-scale sequencing efforts (7, 19–22).

Pathogenic SNPs are enriched at binding sites across ligand types, including protein, nucleic acid, and small molecule binding sites (23–32); yet the relative contributions of distinct binding site types remain unclear. CWMs, in particular, have not been systematically investigated, despite case evidence (33) that their disruption by SNPs can underlie disease.

In Gaucher disease, CWMs have not yet been implicated as a mechanistic factor (34). Responsible for ∼40% of cases worldwide is the L444P variant of the lysosomal acid glucosylceramidase (GCase) encoded by *GBA1* (35). Molecular dynamics (MD) simulations show that L444P, lying in the immunoglobulin-like domain at the interface with the catalytic TIM barrel, destabilizes loops at the active-site entrance, which regulate substrate access, and impairs GCase activation by its facilitator protein, saposin C (34). Therapeutic strategies to stabilize the L444P variant of GCase are emerging for Gaucher disease, yet progress is constrained by limited mechanistic insight, particularly into the role of conserved water molecules (36).

Here, we report the first PDB-wide statistical study linking different binding site types, including CWM sites, to SNP pathogenicity. Using the GenProBiS approach (37–40) we mapped SNPs, classified as benign or pathogenic, onto human protein structures in the PDB and calculated the associations of pathogenic SNPs with individual binding site types. Using MD simulations, we established a causal link between the disruption of a CWM site in the L444P variant of GCase, and the structural changes at the active site entrance. Our results indicate that the loss of a single CWM can explain and extend previously observed effects of this variant in Gaucher disease (34). To support mechanistic studies, pathogenicity prediction, and drug discovery, we provide a freely available dataset mapping SNPs to binding sites, including newly identified CWM sites (*SI Appendix*, Dataset *S1*).

## Results

### SNP pathogenicity inside vs. outside binding sites

We analyzed 4,580 non-redundant SNPs mapped to human protein structures in the PDB, classifying them as within binding sites or on non-binding protein surfaces (Fig. 1*A*). Pathogenic SNPs were more frequent inside than outside binding sites (n = 1,958 vs. 1,257), with a higher overall SNP count (n = 2,466 vs. 2,114) and a greater pathogenic fraction (pf 79.4% vs. 59.5%) (Fig. 1*B* and *C*). SNPs in binding sites of any type were almost three times more likely to be pathogenic than those on non-binding surfaces (odds ratio [OR] = 2.63; 95% confidence interval [CI] 2.31–3.00; *p* = 4.93 × 10^−49^) (Fig. 1*D*).

**Fig. 1.**
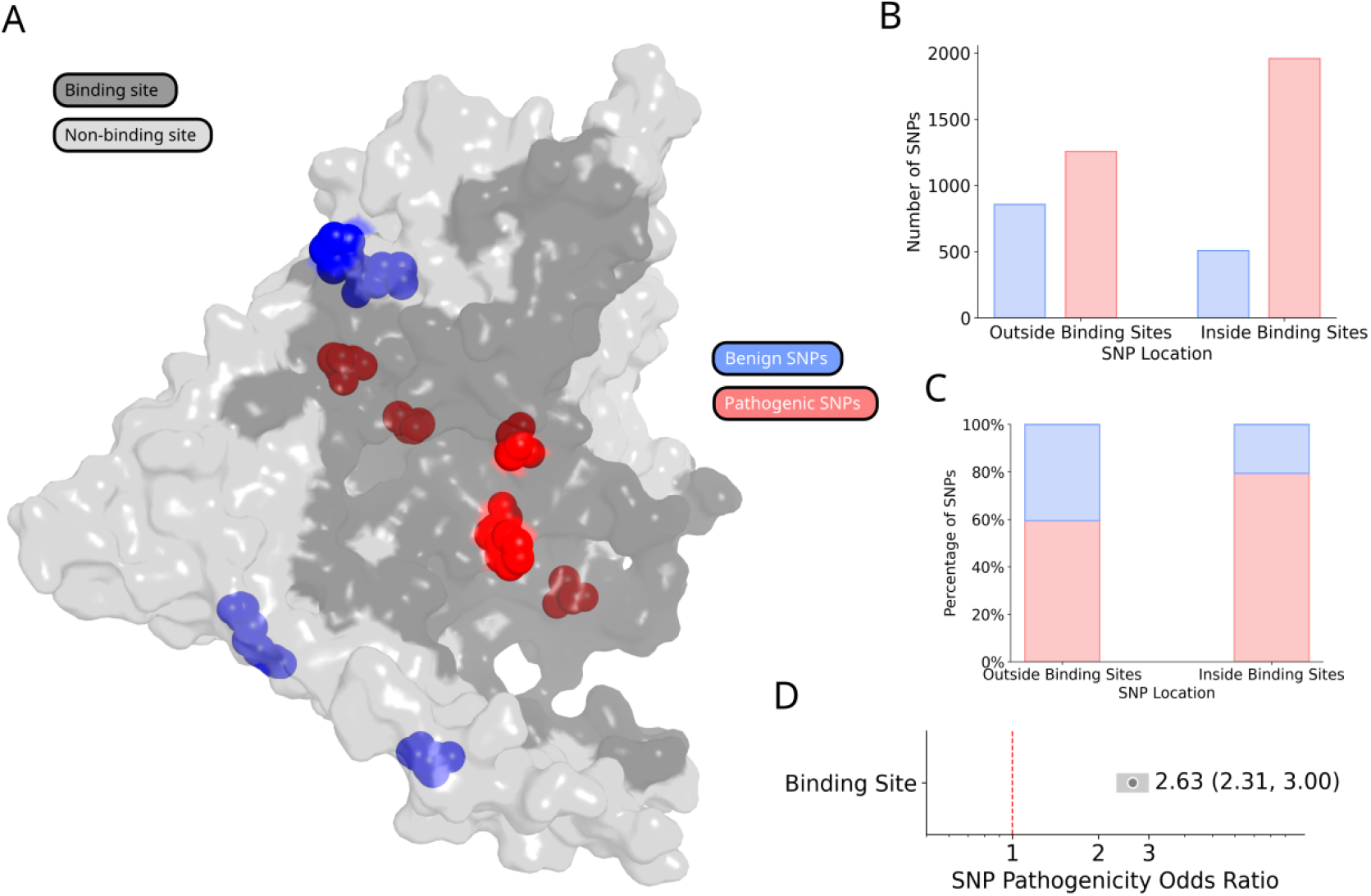
Pathogenic SNPs are enriched in protein binding sites. (*A*) Pathogenic (red) and benign (blue) SNPs mapped to the L-amino acid decarboxylase structure (PDB 3RBF). Binding site surface (dark grey) represents the union of all ligand types. (*B*) Counts of benign (blue) and pathogenic (red) SNPs outside versus inside binding sites. (*C*) Proportion of benign and pathogenic SNPs in each category. (*D*) Odds ratio (OR) and 95% confidence interval (CI) for the association between pathogenic SNPs and binding sites; grey dot, OR; horizontal bar (light grey), CI; dashed vertical red line, OR = 1 (no association).

### Effect of binding site type on SNP pathogenicity

Across binding site types (see examples in Fig. 2*A*), metal ion sites showed the strongest association with pathogenicity (OR = 8.52; CI 4.45–16.32; *p* = 2.66 × 10^−17^; pf = 92.6%; n = 125) (Fig. 2*B* and *C*). Other types were also significantly enriched (*p* < 10^−8^): cofactor (OR = 6.55; CI 4.64–9.25; *p* = 1.49 × 10^−38^; pf = 90.6%; n = 365), compound (OR = 4.10; CI 3.34–5.03; *p* = 2.6 × 10^−17^; pf =85.7%; n = 782), nucleic acid (OR = 3.48; CI 2.14–5.66; *p* = 3.04 × 10^−8^; pf = 83.6%; n = 102), and protein (OR = 2.45; CI 2.12–2.83; *p* = 2.01 × 10^−36^; pf = 78.2%; n = 1,383), whereas glycan sites showed no significant enrichment (OR = 1.09; CI 0.81–1.47; *p* = 0.598; pf = 61.5%; n = 123).

**Fig. 2.**
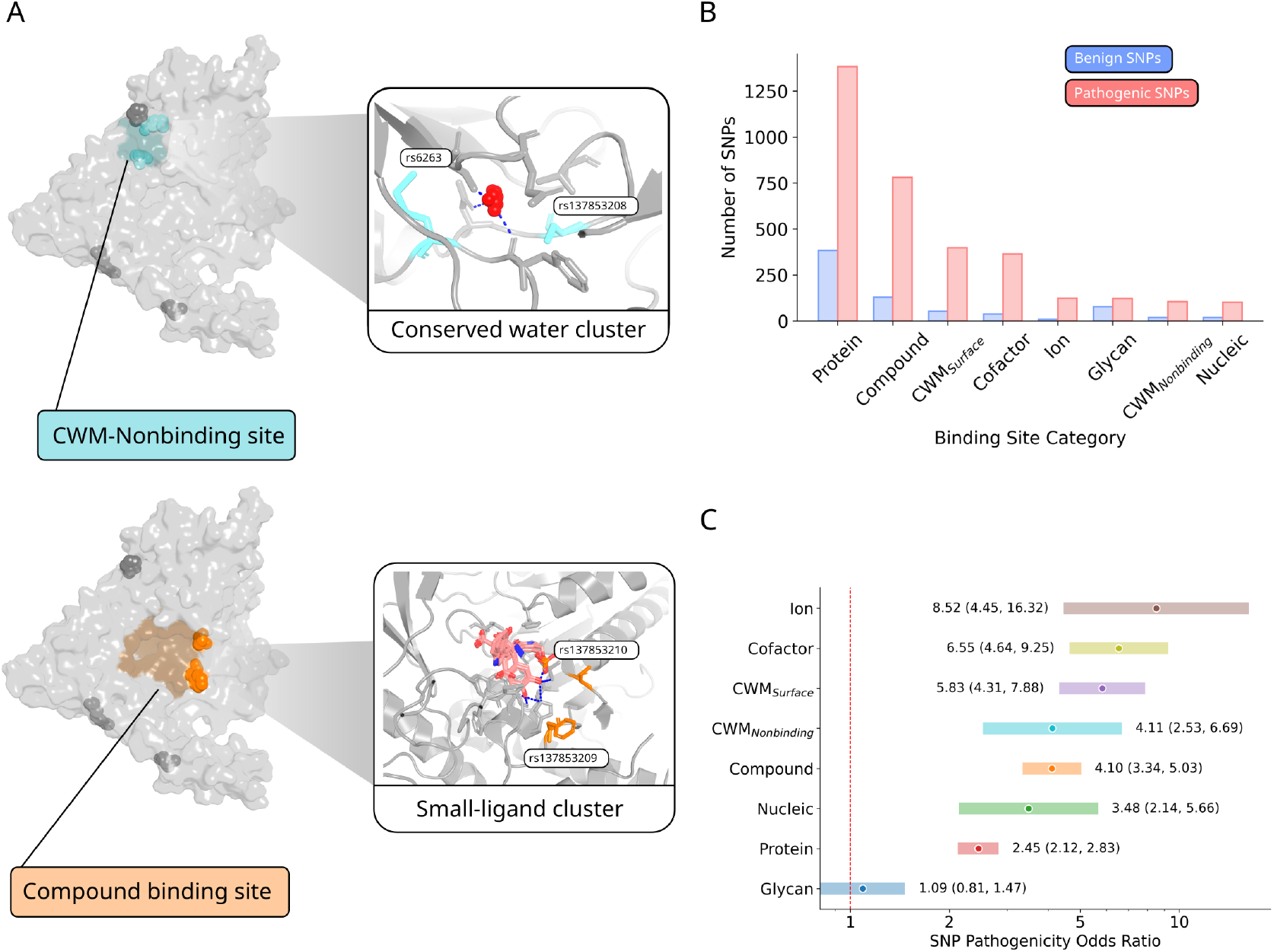
Binding site type–specific effects on SNP pathogenicity. (*A*) SNPs (spheres) mapped to different binding site types (colored surfaces) or to non-binding surface residues (grey) in aromatic-L-amino-acid decarboxylase (*DDC*; PDB 3RBF). Insets show predicted ligands bound to the protein: water molecules (red spheres), compounds (sticks); hydrogen bonds between ligands and protein residues are shown as blue dashed lines; pathogenic SNPs (sticks) are labeled with their rsIDs. (*B*) Counts of benign and pathogenic SNPs by binding site type. (*C*) Odds ratios and 95% CIs for the association between pathogenic SNPs and each binding site type; CWM-surface and CWM-nonbinding site ORs calculated at water conservation level = 0.6; dashed red line, no association.

### Conserved water molecule (CWM) binding sites

To distinguish functional differences, CWM sites were classified as either located anywhere on a protein surface (CWM-surface sites) or restricted to ligand-free surfaces (CWM-nonbinding sites), i.e., outside any other ligand binding site. CWMs within ligand binding sites can mediate ligand recognition or catalysis, whereas those outside binding sites likely have structure-stabilizing roles. CWM-surface sites were strongly associated with pathogenic SNPs (OR = 5.83; CI 4.31–7.88; *p* =4.56 × 10^−41^; pf = 88.3%; n = 399); CWM-nonbinding sites retained substantial enrichment (OR = 4.11; CI 2.53–6.69; *p* = 1.53 × 10^−10^; pf = 84.1%; n = 106) (Fig. 2*B* and *C*).

### Water conservation and evolutionary conservation

In CWM-surface sites, SNP pathogenicity increased with water conservation, rising sharply above conservation level 0.5 and reaching OR = 14.72 at level 1.0 (*p* = 1.05 × 10^−7^), although estimates at the highest level are limited by sample size (Fig. 3*A*). Evolutionary conservation of amino acid residues also correlated with pathogenicity both inside and outside binding sites (Fig. 3*B* and *C*): for residues at the highest conservation level of nine, 94.4% of binding-site SNPs and 81.5% of non-binding SNPs were pathogenic, compared with 73.1% and 51.2% at level zero. Protein–protein sites showed the clearest monotonic increase in pathogenicity with conservation; similar trends were observed in CWM-surface and compound sites, though patterns in other types were less consistent due to small numbers (*SI Appendix*, Fig. *S1*).

**Fig. 3.**
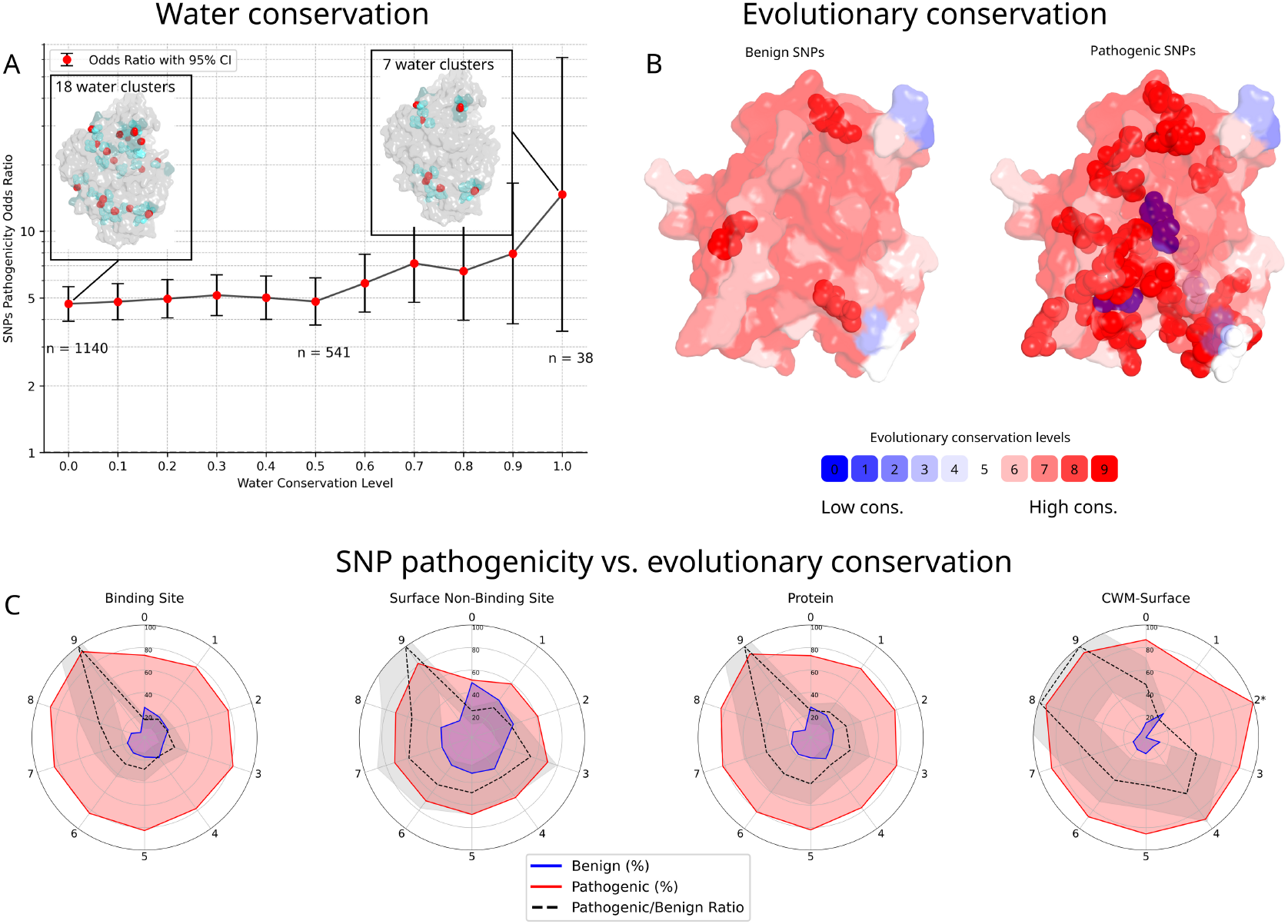
Influence of water and evolutionary conservation on SNP pathogenicity. (*A*) Odds ratios (red dots) and 95% CIs for the association between pathogenic SNPs and CWM sites at varying water conservation thresholds (0.0–1.0). Insets show CWMs (red spheres) in lysosomal acid glucosylceramidase (GCase) structure (*GBA1*; PDB 1OGS) detected at different water conservation thresholds. Number of pathogenic SNPs inside binding sites, n. (*B*) Evolutionary conservation mapped to the surface of the p53 protein (*TP53*; PDB 1TUP) from unconserved (0) to highly conserved (9). (*C*) Radar plots showing the percentage of benign (blue) and pathogenic (red) SNPs, and the normalized pathogenic-to-benign ratio (black dashed line, 95% CI shaded), across evolutionary conservation levels in: all binding sites, non-binding surfaces, protein–protein binding sites, and CWM-surface sites. Asterisks mark conservation levels with missing data, e.g. no benign SNPs in CWM-surface sites at conservation level of two.

### Conserved water molecules and their role in pathogenic remodeling of lysosomal acid glucosylceramidase in Gaucher disease

Pathogenic SNPs were enriched in CWM sites (water conservation ≥0.9) of lysosomal acid glucosylceramidase (GCase), including three variants in CWM-surface sites and twelve in CWM-nonbinding sites (Fig. *4A*; *SI Appendix*, Fig. *S2* and *S3*, Dataset *S1*). The L444P mutation (rs421016; p.Leu483Pro) *mapped to* a CWM-nonbinding site that bridges two β-strands in domain II of GCase via three CWM-mediated hydrogen bonds with Asp443 and Asp445 on the first, and Arg463 on the second β-strand (Fig. *4B*). We therefore hypothesized that the conserved water molecule at this site stabilizes the relative positioning of the opposing β-strands and that the L444P mutation disrupts this stabilization.

**Fig. 4.**
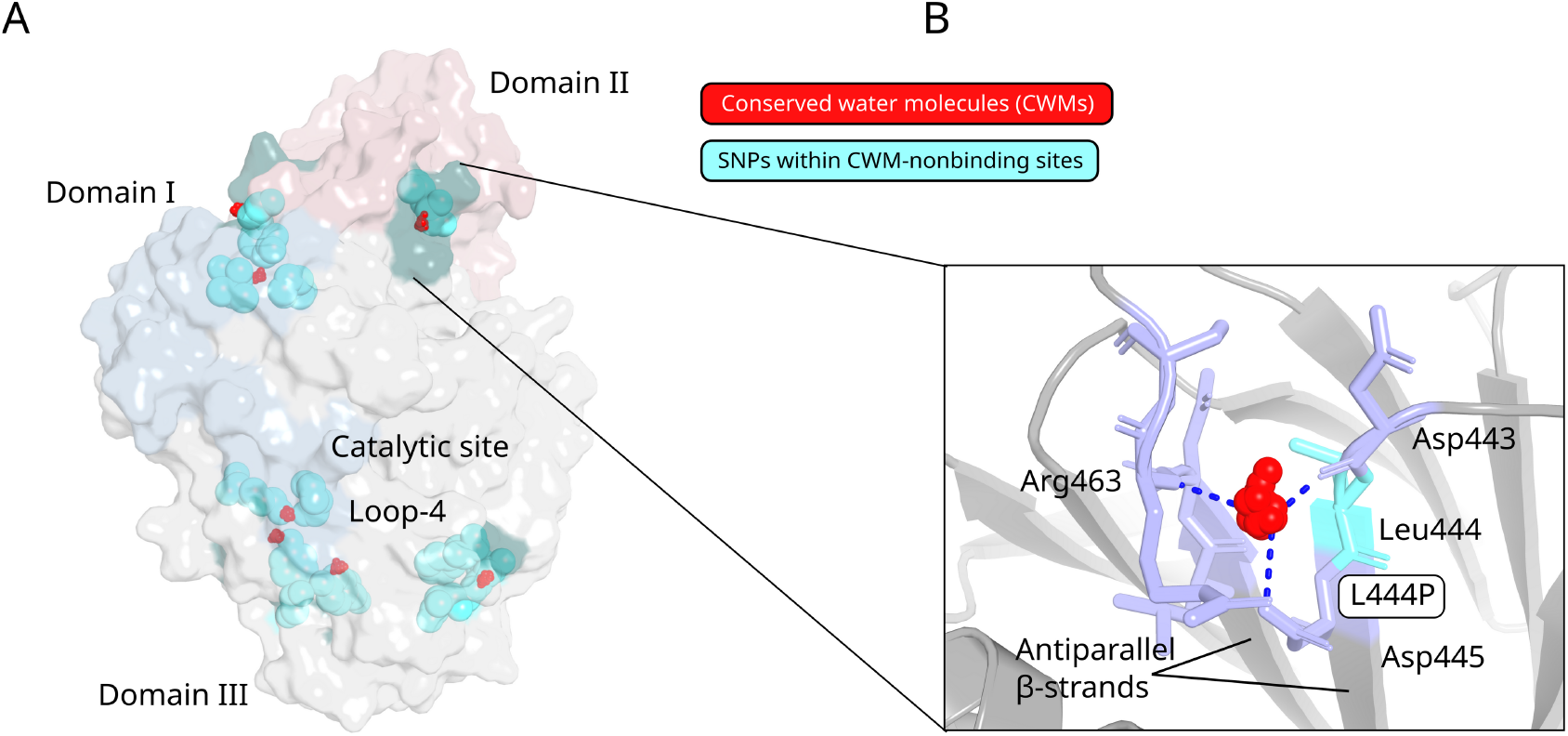
Pathogenic SNP clusters in conserved water molecule binding sites of lysosomal acid glucosylceramidase (GCase). (*A*) Central view of GCase (PDB 1OGS) with CWM-nonbinding sites. CWMs (red spheres, oxygen atoms only) are shown with associated SNP residues (cyan sticks) and other binding site residues (light purple sticks). (*B*) Close-up of a CWM site (light purple sticks) with a cluster of CWMs (red spheres) near the Gaucher disease-causing L444P variant (rs421016; cyan sticks). Hydrogen bonds between conserved waters and protein residues are indicated by blue dashed lines.

To evaluate the structural consequences of CWM disruption, we performed 1-µs molecular dynamics simulations of four systems: wild-type GCase, the L444P variant, a CWM knockout wild-type model, and a rescued L444P variant. In the latter two simulations, the identified conserved water molecule was selectively destabilized or stabilized, respectively (*SI Appendix*, Fig. *S4*).

In the wild-type simulation, the conserved water formed 1,634 bridging hydrogen bonds connecting opposing β-strands. In contrast, both the L444P and CWM knockout simulations exhibited marked reductions in these interactions (1,289 and 1,090, respectively; Fig. *5A* and *B*). Importantly, stabilization of the conserved water molecule in the rescued L444P simulation partially restored β-strand–bridging interactions, increasing their number to 1,332 (see *SI Appendix*, Table *S1*).

**Fig. 5.**
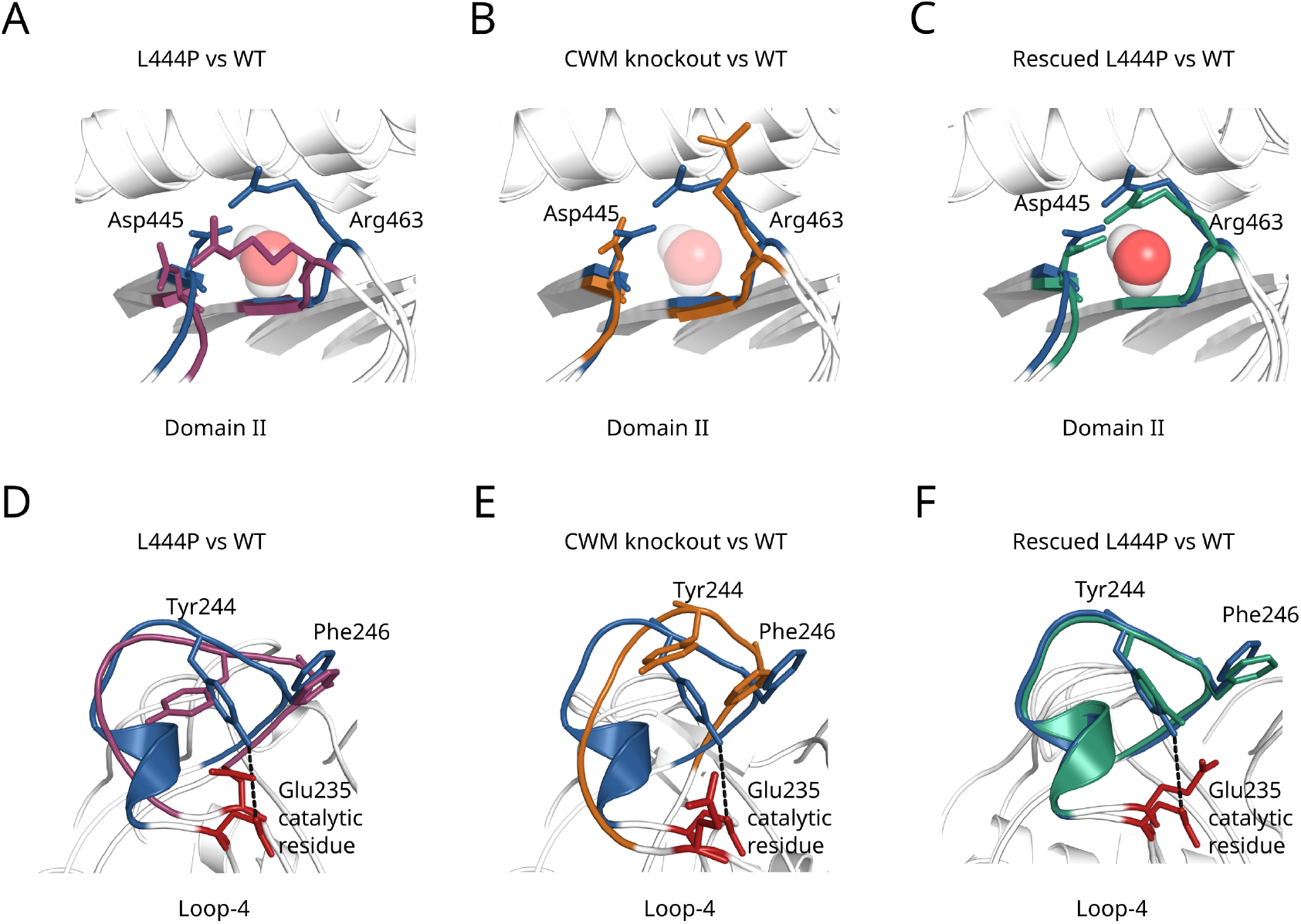
Dynamics of lysosomal acid glucosylceramidase (GCase) variants relative to wild type. (*A–C*) Representative structures of domain II antiparallel β-strands containing the identified conserved water molecule (CWM) for: (*A*) L444P (purple), (*B*) CWM knockout (orange), (*C*) rescued L444P (green), with CWM opacity reflecting hydrogen-bond strength to neighboring residues (higher opacity, more wild-type–like). (*D–F*) Representative structures of loop-4 for: (*D*) L444P, (*E*) CWM knockout, (*F*) rescued L444P. Wild-type structure is dark blue; catalytic Glu235 is red; Tyr244–Glu235 hydrogen bonds in wild-type are dashed lines.

Consistent with loss of β-strand stabilization, both the L444P and CWM knockout simulations exhibited increased separation and broader distance distributions between sidechains of Asp445 and Arg463, which form an inter-β-strand salt bridge in the wild-type simulation (*SI Appendix*, Fig. *S5*). Increased flexibility was also observed in loops surrounding the active site, including loop-4 (residues 237–248; *SI Appendix*, Fig. *S6*).

In wild-type simulation, loop-4 adopted a single dominant conformation with a hydrogen bond between the Tyr244 and catalytic Glu235. In contrast, both the L444P and CWM knockout simulations sampled an additional conformation characterized by rotation of Tyr244 away from Glu235 and partial loss of the short α-helix spanning residues 237–241 (Fig. *5D* and *E*).

Stabilization of the conserved water in the rescued L444P simulation restored local and global dynamics toward wild-type behavior, including recovery of loop-4 conformation and persistent Tyr244–Glu235 hydrogen bonding (Fig. *5C* and *F*; *SI Appendix*, Figs. *S4* to *S6*).

## Discussion

This study presents a proteome-wide structural analysis of pathogenic SNPs across protein binding sites, classified by ligand type and explicitly incorporating conserved water molecules (CWMs). Consistent with prior work (24, 25), SNPs within binding sites are nearly threefold more likely to be pathogenic than those on non-binding protein surfaces, while pathogenic enrichment varies substantially across site classes. Metal ion and cofactor sites show the strongest associations, reflecting stringent physicochemical constraints, whereas protein–protein interfaces exhibit more modest enrichment, consistent with greater mutational tolerance due to larger interface size and partner co-evolution (41, 42). Glycan binding sites show no significant enrichment, likely due to smaller sample size or differences in site definition (43, 44).

A central finding is the strong and independent association between CWMs and pathogenicity. Both CWM sites that occur anywhere on a protein surface (CWM-surface) and CWM sites on otherwise ligand-free surfaces (CWM-nonbinding) are highly enriched for disease-causing variants, and pathogenicity increases with water conservation. These results extend established roles of conserved water in protein stability and ligand recognition (3) and provide, to our knowledge, the first direct association between CWMs and human genetic disease in PDB protein structures.

The causal role of CWMs is demonstrated by the L444P variant of lysosomal acid glucosylceramidase (GCase), a major determinant of Gaucher disease and a Parkinson’s disease risk factor. L444P localizes to a highly conserved CWM-nonbinding site in domain II (immunoglobulin-like domain), where a single CWM bridges two antiparallel β-strands via hydrogen bonds involving residues Asp443–Leu444–Asp445 and Leu461–Asn462–Arg463. Loss of this CWM disrupts hydration-mediated coupling between Asp445 and Arg463, destabilizing local β-sheet packing and increasing structural mobility in this region. Selective removal of the conserved water reproduces the local destabilization and downstream conformational changes observed in the L444P variant, while stabilizing the water suppresses these effects, establishing a direct mechanistic link between CWM loss and pathogenic remodeling.

The CWM destabilization propagates toward the active-site entrance, notably affecting loop-4 (residues 237–248), which contains Tyr244, an aromatic residue lining the catalytic pocket and implicated in substrate recognition (45). Tyr244 lies adjacent to the catalytic dyad Glu235 (acid/base catalyst) and Glu340 (nucleophile), and its orientation modulates access to the active site.

Prior studies show that mutations near the active site can induce conformational rearrangements of Tyr244 and Phe246 (35), consistent with our observation that CWM-mediated destabilization of domain II alters loop dynamics and active-site accessibility without directly perturbing catalytic residues.

The involvement of residues 443–445 in interactions with the facilitator protein saposin C (34) further links our finding of CWM disruption to impaired GCase activation and severe Gaucher disease phenotypes. Consistent with our results, related β-glycosidases from hyperthermophilic organisms have increased numbers of buried water molecules that provide stability by hydrogen bonding to otherwise unsatisfied polar groups within these proteins (45).

Together, our results position CWMs as integral structural determinants whose disruption can drive pathogenic structural remodeling. For Gaucher disease, we highlight conserved hydration sites as mechanistically relevant disease-causing features that may be considered alongside existing approaches aimed at restoring GCase stability and activity (36).

## Materials and Methods

### Detection and classification of human protein binding sites by ligand type

Protein binding sites were identified using the GenProBiS approach (37) (http://genprobis.insilab.org) that implements the ProBiS-ligands method (38). This method scans the entire PDB for local structural similarities and transfers co-crystallized ligands from matched template proteins to the query structure. Ligand clusters, classified into seven types—cofactors, compounds, nucleic acids, proteins, glycans, ions, and conserved water molecules (CWMs)—define distinct binding sites on each query protein. Key steps are:

**Step 1. Filtering for human structures**. PDB protein chains were taxonomically filtered for *Homo sapiens*, yielding 120,628 chains for binding site and ligand detection.

**Step 2. Template identification**. Each human protein chain was used once as a query and compared against ∼320,000 PDB chains using the ProBiS algorithm (40), which detects local surface similarities across whole proteins. Similarity thresholds (Z-scores) were ligand-specific:>2.5 for compounds/cofactors, >3.0 for proteins/nucleic acids, and >2.0 for ions/water (37).

**Step 3. Ligand transposition**. Ligands from templates were mapped to the query via structural alignment if >3 residues (ions, water, small molecules) or >7 residues (proteins, nucleic acids) lay within 4 Å. Ligands causing >10 atoms <1.0 Å steric clashes with the query were discarded.

**Step 4. Binding site detection**. Ligands were clustered by atomic distance using the OPTICS algorithm (46), with artifacts removed via curated nonspecific-binder dataset at http://insilab.org/files/GenProBiS/non-specific.txt; each cluster defined a binding site. Ion/water clusters with <10 members were excluded. Waters with conservation ≥0.6 were classified as CWMs (see below). Binding site residues were defined as those within 5 Å of any ligand in a cluster; glycan sites thus included residues contacting sugar moieties and the covalent attachment site.

### Water conservation at CWM sites

Water conservation quantifies how consistently a bound water molecule is preserved across locally similar protein structures identified by ProBiS. It is calculated as:

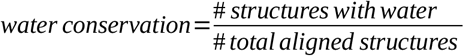

Values range from 0 (unconserved) to 1 (highly conserved) (47). For each SNP within a CWM site, we assigned the highest water conservation level observed among all CWM sites in different protein structures containing that SNP.

### Mapping SNPs to binding sites

SNPs from dbSNP (18) were mapped to protein structures using GenProBiS (37). Only surface SNPs (≤1.1 Å beneath the solvent-accessible surface) were considered; buried SNPs were excluded. SNPs overlapping predicted binding sites were classified by ligand type; overlaps were allowed, so a SNP could belong to multiple binding site categories. ClinVar annotations (48) were used to bin SNPs as pathogenic (pathogenic and likely pathogenic) or benign (benign and likely benign); others were excluded. Evolutionary conservation scores (0–9) for protein surface residues were computed from local structural alignments with ProBiS (39).

### Eliminating SNP redundancy

To avoid overrepresentation from PDB redundancy, SNPs were deduplicated by their reference SNP identifiers (rsIDs), reducing counts from 93,646 to 6,110. SNPs were further collapsed by sequence identity clusters (>30% identity, >20 residues), retaining only those from the chain with the most SNPs in each cluster, yielding 4,580 unique SNPs. For each SNP we recorded: rsID, pathogenicity, binding site type(s), surface non-binding site status, buriedness (such non-binding site SNPs were excluded from analysis but still recorded), PDB/Chain ID, gene name, evolutionary conservation, maximum water conservation level (only for SNPs within CWM sites), amino acid change, and UniProt ID.

### Statistical analysis

Fisher’s exact test was used to compare the frequency of pathogenic SNPs in binding sites vs. non-binding surface regions. Odds ratios were computed for each ligand type and for all sites combined. For CWMs, ‘non-binding surface’ excluded all waters; for other ligand types, SNPs within CWM-nonbinding sites (i.e., CWM sites on otherwise ligand-free surface) were included in the non-binding category. For CWM sites, odds ratios for SNP pathogenicity were computed across water conservation thresholds (0.0–1.0 in 0.1 increments), with the positive set defined as SNPs in CWM sites conserved at or above each threshold, and the negative set as non-binding surface SNPs lacking any CWM site (no conservation threshold applied). A similar framework was applied to evolutionary conservation: SNPs above a conservation threshold were treated as positives, those below as negatives.

### Molecular dynamics simulations of lysosomal acid glucosylceramidase variants

Molecular dynamics simulations were performed on human lysosomal acid glucosylceramidase (GCase) using the crystal structure PDB 1OGS (chain A). Four systems were simulated: (i) wild-type GCase, (ii) GCase with the L444P mutation, (iii) conserved water molecule–knockout (“CWM knockout”) variant of wild-type GCase, and (iv) a CWM-stabilized L444P variant (“rescued L444P”). The L444P mutation was introduced using PDBFixer in OpenMM (49). All systems were solvated in explicit TIP3P-FB water with a 1.0 nm padding and neutralized at 0.15 M ionic strength using K^+^ and Cl^—^ ions. Simulations employed the AMBER14 force field and particle mesh Ewald electrostatics with a 1.0 nm nonbonded cutoff.

In the CWM knockout simulation, electrostatic interactions between bulk water and two protein backbone amide hydrogens (Asp445 and Arg463) were selectively removed using a custom nonbonded force in OpenMM. Electrostatic terms between these protein atoms and all water atoms were set to zero, while van der Waals interactions and all other protein electrostatics were preserved, thereby destabilizing the conserved water site without otherwise perturbing the wild-type protein structure.

In the rescued L444P simulation, electrostatic interactions anchoring the conserved water molecule at its binding site were selectively strengthened. This was achieved by scaling the electrostatic terms between the Asp443 backbone carbonyl oxygen and bulk water hydrogens, and between the Arg463 amide hydrogen and bulk water oxygens, by a factor of 1.1. To permit exchange of the conserved water molecule with bulk solvent, as observed in the wild-type simulation, this scaling was applied intermittently in 100,000-step cycles (25,000 steps on, 75,000 steps off), using the L444P structure as the starting model.

All systems were energy-minimized (5,000 steps), heated to 310.15 K over 10 ns under NPT conditions at 1 bar using a Langevin integrator, and simulated for 1 μs. The first 100 ns were discarded as equilibration, and the remaining 900 ns were used for analysis.

Trajectory analysis was performed using MDAnalysis (50). Hydrogen bonds between water molecules and protein residues were identified using geometric criteria implemented in HydrogenBondAnalysis and evaluated at regular intervals along the trajectory. Conformational clustering was performed on superposed trajectories aligned on protein Cα atoms using an automated k-means procedure with silhouette scoring to select the optimal number of clusters; representative structures were defined as cluster medoids.

## Data Availability

Data to generate SNP variants annotated by their pathogenicity and involvement in CWM sites and specific ligand-binding sites for protein structures in the PDB and molecular dynamics trajectories of GCase variants are freely available at http://insilab.org/files/share/patho-snps-water-ligand-dataset.zip and at http://insilab.org/files/share/gcase-variants-md.zip.

## Code Availability

Software developed for the statistical analysis and visualization of the SNPs mapped to ligand-binding sites and CWM sites is freely available at https://gitlab.com/janezkonc/patho-snp-water-ligand. Molecular dynamics scripts are available at https://gitlab.com/janezkonc/gaucher-md.

## Funding

This work was supported by the Slovenian Research and Innovation Agency (ARIS), Research program P4-0220 (T.K.) and project grants N1-0142 (J.K.), N1-0209 (D.J.), J1-4414 (D.J.), and L7-8269 (J.K.).

## Author Contributions

J.K., T.K. and D.J. conceived and designed the study. J.K. and K.R. performed the computational experiments and collected the data. J.K. and K.R. conducted data analysis and interpretation. J.K. prepared figures and tables. All authors wrote the main manuscript text. All authors reviewed and approved the final manuscript.

## Competing Interests

The authors declare no competing interests.

## Supplementary Information

**Supplementary Fig. 1: Evolutionary conservation and SNP pathogenicity across binding site types**. Radar plots showing the percentage of benign (blue) and pathogenic (red) SNPs, and the normalized pathogenic-to-benign ratio (black dashed line, 95% CI shaded), across evolutionary conservation levels for all binding site types. Asterisks indicate conservation levels with missing data (e.g., no benign SNPs in CWM-surface sites at level 2).

**Supplementary Fig. S2. Pathogenic SNPs within CWM-surface and CWM-nonbinding sites (water conservation ≥ 0.9) mapped to the lysosomal acid glucosylceramidase (GCase) structure (GBA1; PDB 1OGS)**.

**Supplementary Fig. S3. Lysosomal acid glucosylceramidase (GCase) with close-ups of CWM-nonbinding sites**. Conserved water molecules (red spheres, oxygen atoms only) are shown with associated SNP residues (cyan sticks) and other binding site residues (light purple sticks). Hydrogen bonds between conserved waters and protein residues are indicated by blue dashed lines.

**Supplementary Fig. S4. Cα RMSD of GCase variants as a function of simulation time**.

**Supplementary Fig. S5. Distance between Asp445 and Arg463 side chains in GCase variants as a function of simulation time**.

**Supplementary Fig. S6. Residue-wise root mean square fluctuation (RMSF) profiles in GCase variants**.

**Supplementary Table S1. Counts and relative strengths of bridging hydrogen bonds for the conserved water molecule near L444P in GCase variants**.

**Supplementary Dataset S1. Non-redundant SNPs mapped to conserved water and ligand-binding sites**. A total of 5,312 SNPs are annotated with pathogenicity, ligand-binding site classifications, evolutionary conservation, and structural context. Analysis was performed on the 4,580 SNPs located in binding sites or on the non-binding protein surface; the remaining 732 buried SNPs (recognized by non-binding = 1 and non-binding surface = 0) were excluded. Data are organized in separate sheets according to water conservation thresholds (0.0-1.0).

## Notes

### Competing Interest Statement

The authors have declared no competing interest.

