## Supporting Information for "Conserved water molecules as structural ligands modulating pathogenic variation in human protein binding sites"

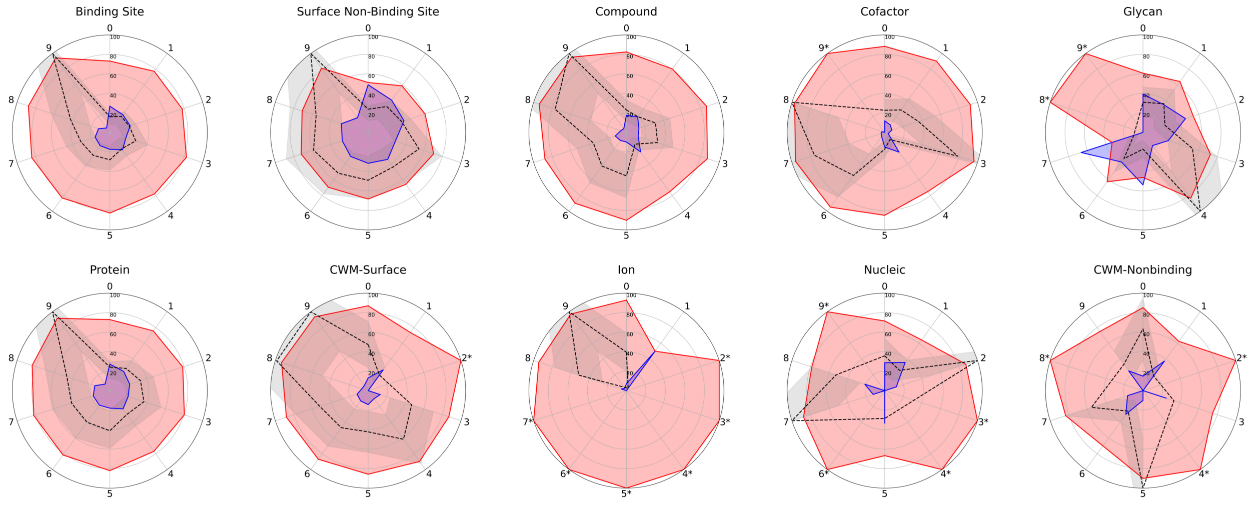


**Fig. S1. Evolutionary conservation and SNP pathogenicity across binding site types.** Radar plots showing the percentage of benign (blue) and pathogenic (red) SNPs, and the normalized pathogenic-to-benign ratio (black dashed line, 95% CI shaded), across evolutionary conservation levels for all binding site types. Asterisks indicate conservation levels with missing data (e.g., no benign SNPs in CWM-surface sites at level 2).

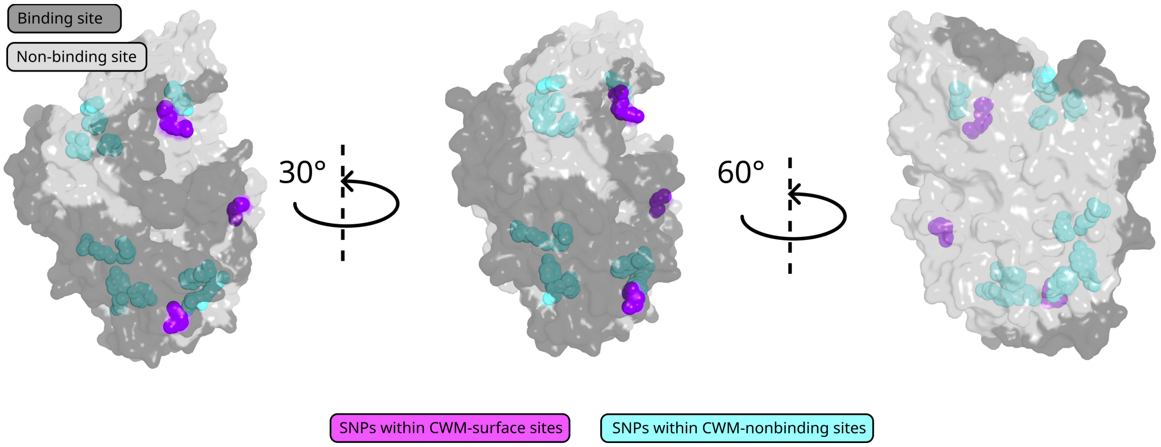


**Fig. S2. Pathogenic SNPs within CWM-surface and CWM-nonbinding sites (water conservation ≥ 0.9) mapped to the lysosomal acid glucosylceramidase (GCase) structure (GBA1; PDB 1OGS).**

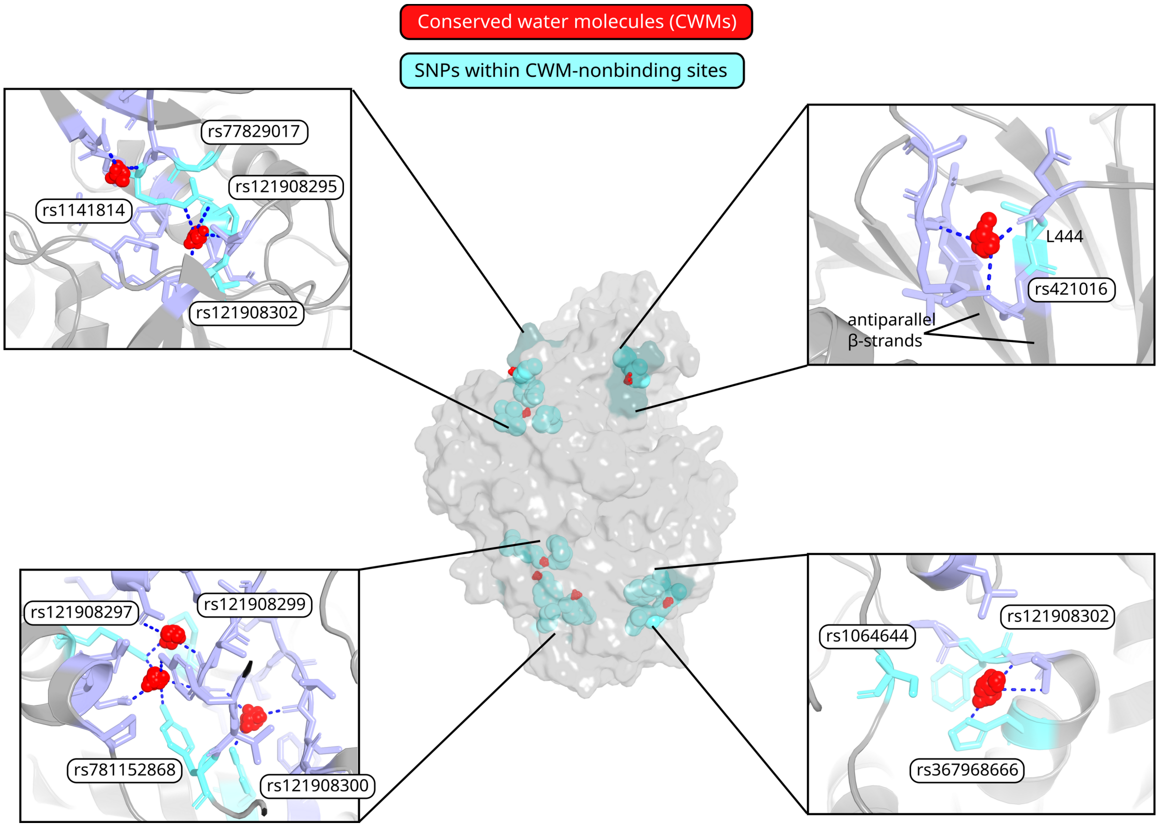


**Fig. S3. Lysosomal acid glucosylceramidase (GCase) with close-ups of CWM-nonbinding sites.** Conserved water molecules (red spheres, oxygen atoms only) are shown with associated SNP residues (cyan sticks) and other binding site residues (light purple sticks). Hydrogen bonds between conserved waters and protein residues are indicated by blue dashed lines.

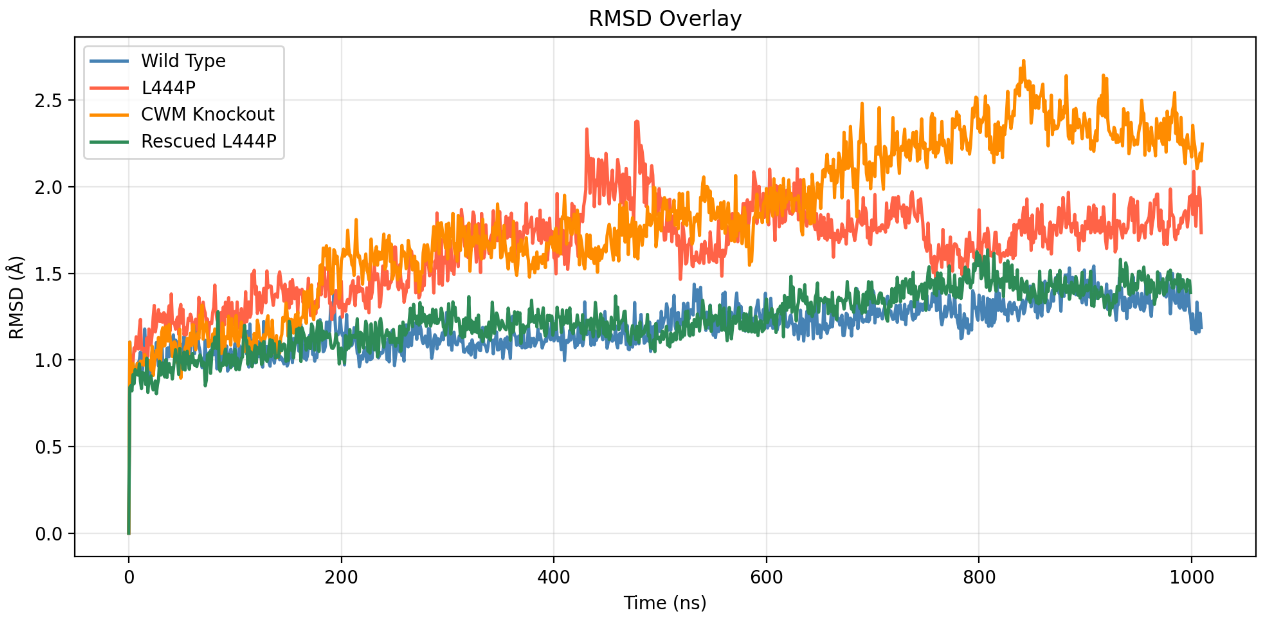


**Fig. S4. Cα RMSD of GCase variants as a function of simulation time.**

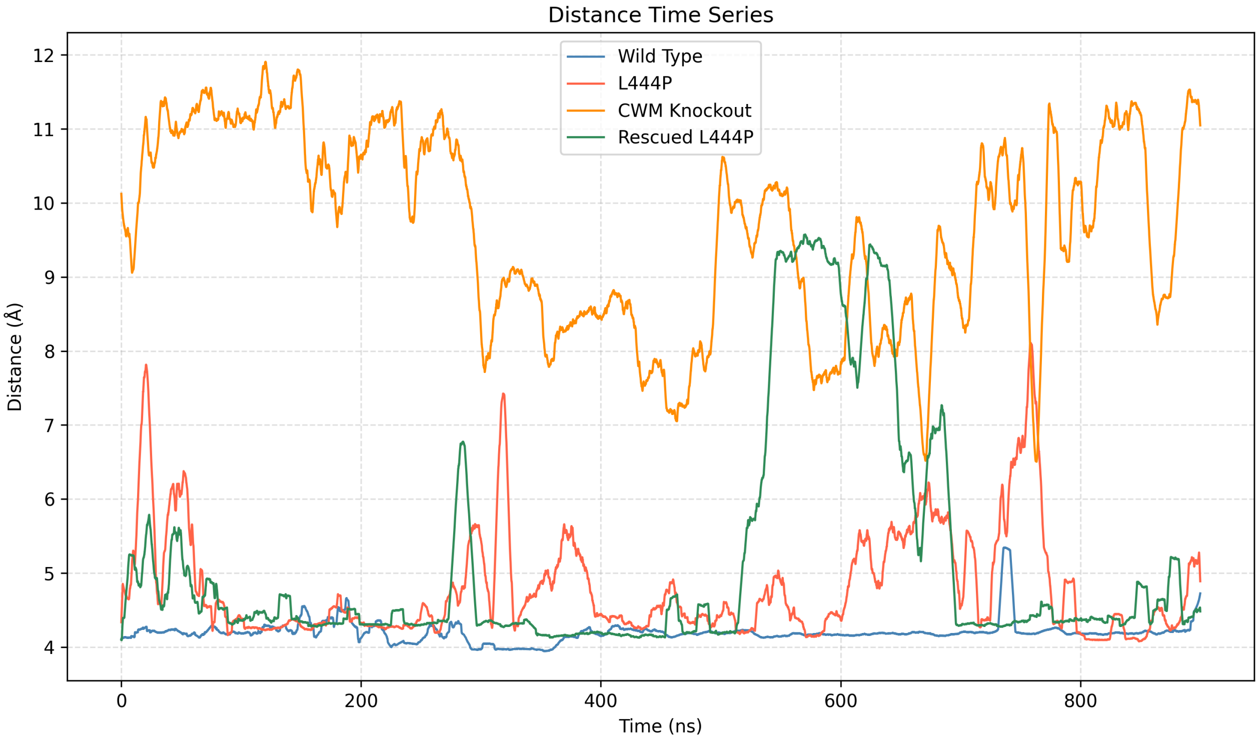


**Fig. S5. Distance between Asp445 and Arg463 side chains in GCase variants as a function of simulation time.**

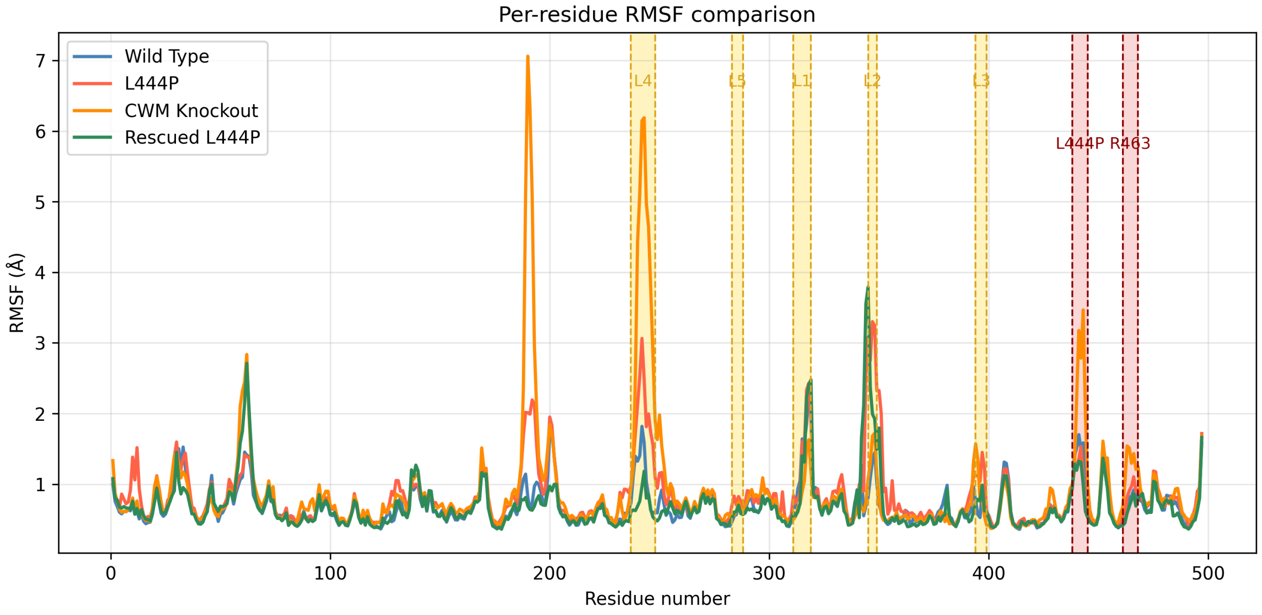


**Fig. S6. Residue-wise root mean square fluctuation (RMSF) profiles in GCase variants.**

**Table S1. Counts and relative strengths of bridging hydrogen bonds for the conserved water molecule near L444P in GCase variants.**

| | Count of bridging CWM hydrogen bonds[1] | Relative strength of bridging CWM hydrogen bonds vs. wild-type[2] |
|---|---|---|
| **Wild Type** | 1634 | 1.0 |
| **L444P** | 1289 | 0.78 |
| **CWM Knockout** | 1090 | 0.67 |
| **Rescued L444P** | 1332 | 0.83 |

[1] A bridging hydrogen bond is defined as a hydrogen bond formed by a water molecule that forms at least two hydrogen bonds, the first one with protein residues 443, 444 or 445, and the second hydrogen bond with residues 461, 462 or 463. These hydrogen bonds were sampled at 100-frame intervals from the last 900 ns of each molecular dynamics simulation.

[2] The relative strength of CWM–protein hydrogen bonds was defined as the ratio of the number of bridging CWM hydrogen bonds in each GCase variant to that in the wild-type GCase, computed over the final 900 ns of each MD simulation. In the CWM knockout simulation, hydrogen bonds involving (“ARG”, 463, “A”, “H”) or (“ASP”, 445, “A”, “H”) were assigned an electrostatic scaling factor of 0.0. In the rescued L444P simulation, hydrogen bonds involving (“ARG”, 463, “A”, “H”) or (“ASP”, 443, “A”, “O”) that occurred during electrostatics-on frames were scaled by a factor of 1.1. Of 1,332 total bridging CWM hydrogen bonds, 289 occurred in electrostatics-on frames, yielding a relative H-bond strength of $(1332 - 289 + (1.1 \times 289)) / 1634$.

**Dataset S1 (separate file). Non-redundant SNPs mapped to conserved water and ligand-binding sites.** A total of 5,312 SNPs are annotated with pathogenicity, ligand-binding site classifications, evolutionary conservation, and structural context. Analysis was performed on the 4,580 SNPs located in binding sites or on the non-binding protein surface; the remaining 732 buried SNPs (recognized by non-binding = 1 and non-binding surface = 0) were excluded. Data are organized in separate sheets according to water conservation thresholds (0.0-1.0).
